# Fitting the Erlang distribution to cancer incidence by age may predict key carcinogenic events

**DOI:** 10.1101/060970

**Authors:** Aleksey V. Belikov

## Abstract

Cancer is the second-leading cause of death worldwide, after cardiovascular diseases. Cancers arise from various cells and organs at different ages and develop at different rates. However, the reasons for this variation in the cancer progression rate and the age of onset are poorly understood. Especially puzzling is the late-life decrease in cancer incidence, which cannot be explained by previously proposed power law or exponential growth equations. By using the latest publicly available USA cancer incidence statistics, comprised of 20 million cancer cases documented over 14 years, I show that cancer incidence by age closely follows the Erlang probability distribution (R^2^=0.9543-0.9999), which is a special case of the gamma distribution. The Erlang distribution describes the probability *y* of *k* independent random events occurring by the time *x*, but not earlier or later, with each event happening on average every *b* time intervals. This fits well with the multiple-hit hypothesis, and potentially allows to predict the number *k* of key carcinogenic events and the average time interval *b* between them, for each cancer type. Moreover, the amplitude parameter *A* likely predicts maximal populational susceptibility to a given type of cancer. These parameters are estimated for 20 most common cancer types, and provide clues for further research on cancer development.

## Introduction

The value of cancer incidence and mortality curves for inferring information about the underlying carcinogenic processes has long been recognized^1^. It has been the basis for the influential multi-hit hypothesis of cancer development, which proposed that cancer appears after seven consecutive mutations^2–4^. That prediction was based on the assumption that cancer mortality increases proportionally to the sixth power of age. However, already at that time it was known that many cancers display deceleration of mortality growth at advanced age, which could not be explained by the power law. Many complicated equations based on multiple assumptions and empirically estimated parameters have since been proposed, attempting to model the limited growth of cancerous cells^5–7^. However, current data unequivocally show that cancer incidence not only ceases to increase with age but, for at least some cancers, decreases^8^. This behavior cannot be explained by any growth equations, and has been puzzling biologists and clinicians for considerable time. Here I propose that cancer incidence by age is in fact a statistical distribution of probabilities that a required number of mutations is accumulated by the given age. Of 17 tested continuous distributions, the best fit is observed for the Erlang distribution, which is a special case of the gamma distribution with an integer shape parameter. Notably, the Erlang distribution describes the probability of several independent random events occurring by the given time. This takes the multiple-hit hypothesis on a new level, and potentially allows to predict the number of key carcinogenic events and the average time interval between them, for each cancer type.

**Figure 1.**
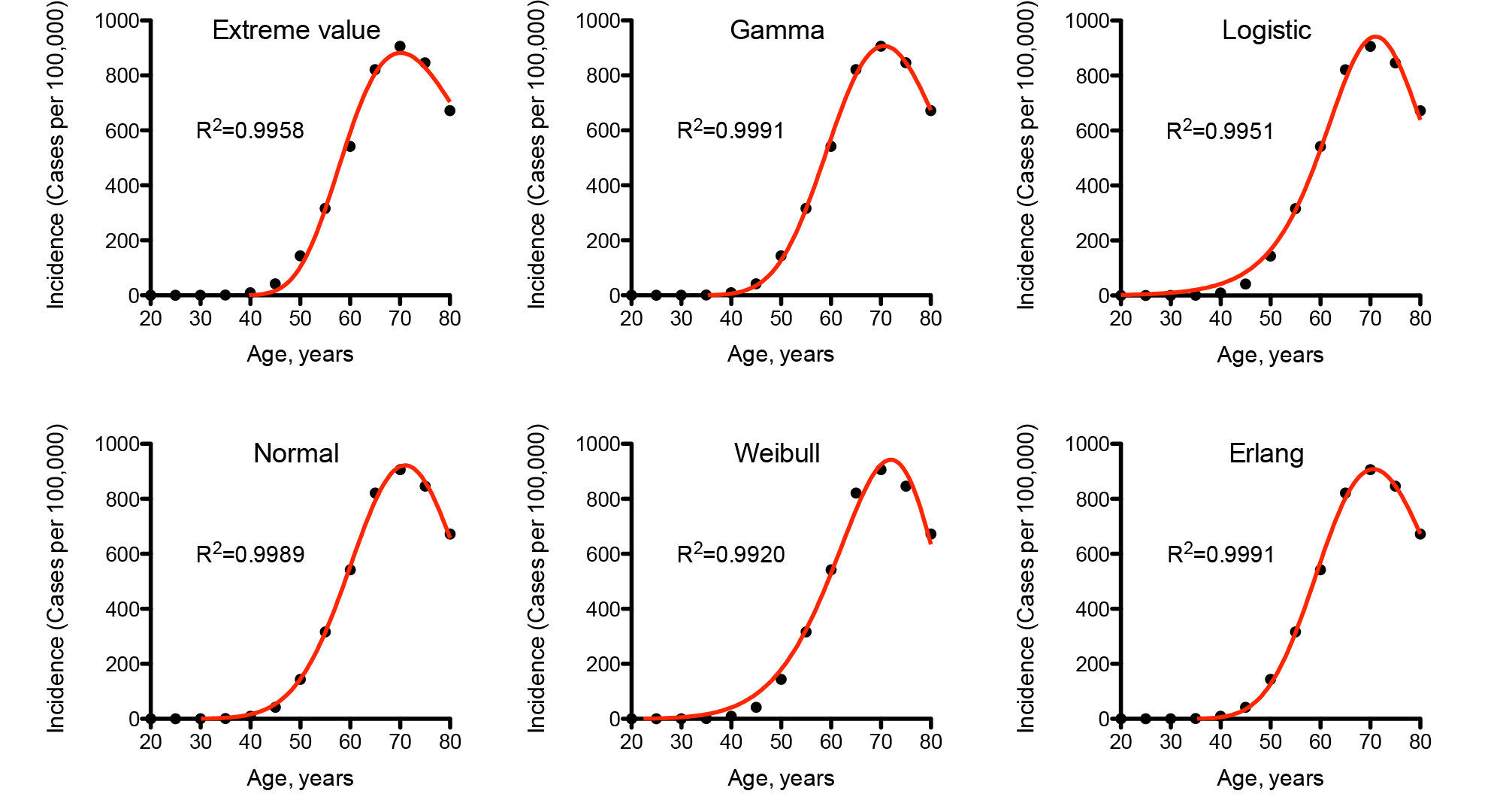
Comparison of different statistical distributions with actual distribution of prostate cancer incidence by age. Dots indicate actual data for 5-year age intervals, curves indicate regressions. The starting age of each age group is indicated. The fitting procedure was identical for all distributions. Prostate cancer was selected due to both the highest incidence and the highly efficient screening procedure.

## Results

To test the probability hypothesis, the latest publicly available USA cancer incidence data were downloaded from the CDC WONDER database (see Methods for details). The probability density functions for the general forms of the following continuous probability distributions were tested for fit with least squares nonweighted nonlinear regression analysis: beta, beta prime, Cauchy, extreme value, Fisher F, gamma, Gompertz, chi-square, Levy, logistic, Maxwell, normal, Rayleigh, Student t, Wald and Weibull. Only the extreme value, gamma, logistic, normal and Weibull distributions provided acceptable fit for most of cancer types, with gamma providing the best fit (Figure 1). The special case of the gamma distribution with integer shape parameter – the Erlang distribution - was also tested and provided the fit identical to the gamma distribution (Figure 1). Whilst the normal distribution also provided a very good fit, not much information can be inferred from it, except of the age of maximal cancer incidence. On the contrary, the parameters of the Erlang distribution can be interpreted in a way to get insights into the carcinogenesis process.
I propose that the shape parameter ***k*** of the Erlang distribution indicates the number of carcinogenic events that need to occur in order for a cancer to develop to a stage that can be detected during clinical screening or by a patient himself. The scale parameter ***b*** indicates the average time interval (in years) between such events. Finally, the amplitude parameter ***A*** divided by 1000 estimates the maximal susceptibility (in percent) of a given population to a given type of cancer. This is because the area under the probability density function curve is always unity, the maximal area under the cancer incidence curve is 100000, and ***A*** is used to convert probabilities into incidence, which is measured in cases per 100000 people.

**Figure 2.**
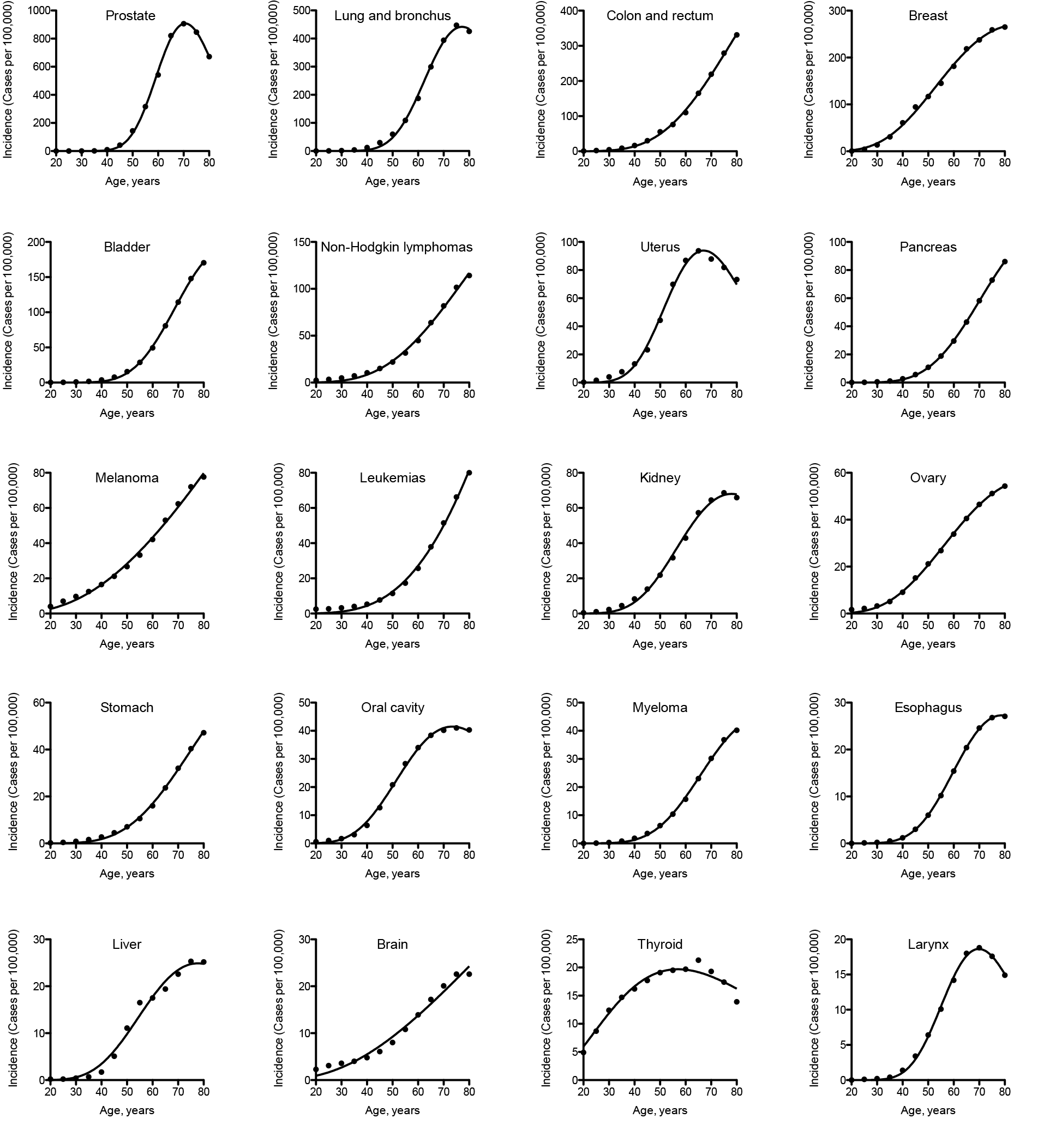
The Erlang distribution approximates cancer incidence by age for 20 most prevalent cancer types. Dots indicate actual data for 5-year age intervals, curves indicate the Erlang regression. The starting age of each age group is indicated.

To obtain these parameter values, the Erlang distribution was fit individually to the data for each of the 20 most prevalent cancer types (Figure 2, Table 1). The goodness of fit varied from 0.9543, for thyroid cancer, to 0.9999, for pancreatic and esophageal cancers, with the average of 0.9943. The predicted number of carcinogenic events varied from 4, for melanoma, brain and thyroid cancers, to 38, for prostate cancer. The predicted average time between the events varied from 2 years, for prostate cancer, to 80 years, for melanoma. The predicted maximal populational susceptibility varied from 1%, for esophageal and larynx cancers, to 100%, for melanoma. Overall, the data predict high heterogeneity in carcinogenesis patterns.

**Table 1.** Estimated carcinogenesis parameters for 20 most prevalent cancer types. The parameters are determined for the Erlang distribution that approximates actual cancer incidence by age.

| Parameter | $k$ | $b$ | $A/1000$ | $R^2$ |
| --- | --- | --- | --- | --- |
| Proposed meaning | Number of carcinogenic events ± SE | Average time between events, years ± SE | Maximal populational susceptibility, % ± SE | Goodness of fit |
| Prostate | 38±1 | 1.91±0.00 | 26.48±0.19 | 0.9991 |
| Lung and bronchus | 28±2 | 2.86±0.01 | 16.51±0.26 | 0.9979 |
| Colon and rectum | 10±1 | 12.92±0.16 | 57.09±3.12 | 0.9990 |
| Breast | 8±1 | 12.13±0.12 | 21.95±0.53 | 0.9981 |
| Bladder | 20±1 | 4.69±0.02 | 9.81±0.18 | 0.9994 |
| Non-Hodgkin lymphomas | 8±1 | 17.95±0.53 | 26.09±3.06 | 0.9962 |
| Uterus | 19±1 | 3.73±0.02 | 3.74±0.06 | 0.9949 |
| Pancreas | 14±1 | 7.46±0.02 | 7.14±0.07 | 0.9999 |
| Melanoma | 4±1 | 80.18±7.18 | 104.81±29.41 | 0.9956 |
| Leukemias | 7±2 | 31.81±1.85 | 91.09±24.79 | 0.9963 |
| Kidney | 14±1 | 6.00±0.05 | 3.71±0.08 | 0.9967 |
| Ovary | 8±1 | 12.90±0.10 | 4.95±0.11 | 0.9990 |
| Stomach | 11±1 | 10.86±0.15 | 6.33±0.36 | 0.9985 |
| Oral cavity | 12±1 | 6.68±0.04 | 2.32±0.03 | 0.9983 |
| Myeloma | 15±1 | 6.41±0.04 | 2.69±0.06 | 0.9991 |
| Esophagus | 18±0 | 4.63±0.01 | 1.31±0.01 | 0.9999 |
| Liver | 12±2 | 7.10±0.13 | 1.47±0.07 | 0.9853 |
| Brain | 4±1 | 67.45±10.45 | 19.26±8.89 | 0.9791 |
| Thyroid | 4±1 | 19.01±0.44 | 1.67±0.05 | 0.9543 |
| Larynx | 23±1 | 3.17±0.01 | 0.70±0.01 | 0.9987 |

## Discussion

The progression from one carcinogenesis stage to the other is usually assumed to be mediated by “driver” mutations in crucial genes, which give mutated cell growth advantage, apoptosis resistance or other oncogenic properties, as opposed to inconsequential “passenger” mutations^9^. Many algorithms have been suggested for identification of driver mutations^10^, indicating that no universally accepted criteria exist. Moreover, whilst hundreds of potential driver mutations have been identified in various tumors, they need not be all present in the same tumor specimen, as many of them are redundant or even mutually exclusive, e.g. when the affected proteins are components of the same signaling pathway^11^. Thus, each tumor is expected to have only a sample of all possible driver mutations. Another aspect to consider is that while one mutation is usually sufficient to activate an oncogene, two mutations are typically required to inactivate both alleles of a tumor suppressor gene. Therefore, the number of carcinogenetic events predicted by the Erlang distribution should be translated not into the number of mutated genes, but rather into the number of mutations.

When cancer drivers are searched for in tumor genomes, most studies focus on nonsynonymous point mutations^12^. This gives relatively low numbers of driver mutations, in the range from one to eight (Fig3 in Ref^12^). However, it has been recently shown that synonymous^13^ and noncoding^14^ mutations also can act as carcinogenesis drivers. Moreover, there are many more types of genetic alterations that can possibly contribute to cancer progression. They include indels^15^, homozygous deletions^16^, inversions^17^, tandem duplications^18^, amplifications^19^, intra- and inter-chromosomal translocations^20^ (often resulting in gene fusions^21^), as well as chromosomal arm-level and whole-level copy-number alterations^22^, and chromothripsis^23^. Additionally, epigenetic alterations (epimutations) are a whole new level of potential cancer drivers^24,25^.

It is likely that many of these alterations contribute to progression of each cancer type. Moreover, different cancer types and subtypes require different proportions of these alterations^26^, e.g. some cancers are driven mostly by point mutations, some by amplifications, yet some by gene fusions. Interestingly, the total number of important alterations per tumor ranged from 0 to 40 (Fig2c in Ref^26^), which corresponds to the range of event numbers predicted by the Erlang distribution. Therefore, the number of carcinogenic events per tumor predicted by the current theory is most likely the sum of alterations of several different types. Astonishingly, the recent massive omics study of 333 primary prostate carcinomas by The Cancer Genome Atlas Research Network has found only a single or no alterations in up to 26% of tumor samples^27^. In extreme case, this may mean that the true nature of carcinogenesis drivers is still not known.

Most data that were used in this study represent combined cancer cases, e.g. acute and chronic, lymphocytic, myeloid and monocytic leukemias were combined into Leukemias. The resulting curve is necessary different in shape, position and amplitude from the curves of individual leukemia subtypes. Hence, the estimated parameters are also different, and reflect only the average. When the exact number of carcinogenic alterations is required, it is necessary to analyze the data for a particular cancer subtype, and also for a particular gender and race. Such data are readily accessible at the CDC WONDER portal.

Another factor that influences the results is the stage at which cancer is diagnosed. Cancer types that are diagnosed at early stages, e.g. due to highly developed screening programs, will likely undergo fewer carcinogenic events by the time of first diagnosis than cancers that are difficult to diagnose early. Thus, the current theory predicts the average number of carcinogenic events that happen by the time of diagnosis, and not by the time of full cancer development.

Overall, the theory and methodology presented here allow to generate testable predictions about the carcinogenesis process in any cancer subtype for which reliable incidence statistics is available. Thus, they may help to define the subtype-specific cancer drivers, by providing numerical reference points. Also, the estimated maximal populational susceptibility may help to predict the allele frequencies of driver genes. Finally, these findings provide additional support to the multiple-hit theory of carcinogenesis.

## Methods

### Data acquisition

United States Cancer Statistics Public Information Data: Incidence 1999 - 2012 were downloaded via Centers for Disease Control and Prevention Wide-ranging OnLine Data for Epidemiologic Research (CDC WONDER) online database (http://wonder.cdc.gov/cancer-v2012.HTML. The United States Cancer Statistics (USCS) are the official federal statistics on cancer incidence from registries having high-quality data for 50 states and the District of Columbia. Data are provided by The Centers for Disease Control and Prevention National Program of Cancer Registries (NPCR) and The National Cancer Institute Surveillance, Epidemiology and End Results (SEER) program. Results were grouped by 5-year Age Groups and Crude Rates were selected as output. All other parameters were left at default settings. Then the data were downloaded separately for each cancer type, upon its selection in the Leading Cancer Sites tab.

### Data selection and analysis

For analysis, the data were imported into GraphPad Prism 5. The following age groups were selected: “20-24 years”, “25-29 years”, “30-34 years”, “35-39 years”, “40-44 years”, “45-49 years”, “50-54 years”, “55-59 years”, “60-64 years “, “65-69 years”, “7074 years”, “75-79 years” and “80-84 years”. Prior age groups were excluded due to unreliably low incidence rates, and “85+ years” was excluded due to the undefined age interval. Data were analyzed with Nonlinear regression. The following User-defined equations were created for the statistical distributions:

*Extreme value:*

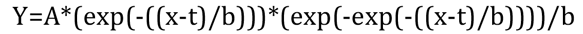

*Gamma:*

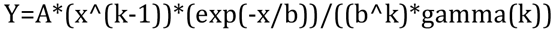

*Logistic:*

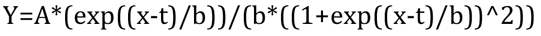

*Normal:*

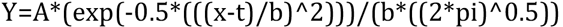

*Weibull:*

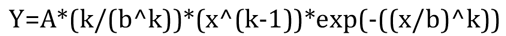

The parameter *A* was constrained to “Must be between zero and 100000.0”, parameter *t* to “Must be between zero and 150.0”, parameters *b* and *k* to “Must be greater than 0.0”. “Initial values, to be fit” for all parameters were set to 1.0. All other settings were left by default, e.g. Least squares fit and No weighting.
For the Erlang distribution, the parameter *k* for each cancer type was estimated by the Gamma regression, rounded to the nearest integer and used as “Constant equal to” in the second round of the Gamma regression, which provided the final results.

